# Preadaptation of pandemic GII.4 noroviruses in hidden virus reservoirs years before emergence

**DOI:** 10.1101/658765

**Authors:** Christopher Ruis, Lisa C. Lindesmith, Michael L. Mallory, Paul D. Brewer-Jensen, Josephine M. Bryant, Veronica Costantini, Christopher Monit, Jan Vinjé, Ralph S. Baric, Richard A. Goldstein, Judith Breuer

## Abstract

The control of pandemic pathogens depends on early prediction of pandemic variants and, more generally, understanding origins of such variants and factors that drive their global spread. This is especially important for GII.4 norovirus, where vaccines under development offer promise to prevent hundreds of millions of annual gastroenteritis cases. Previous studies have suggested that new GII.4 pandemic viruses evolve from previous pandemic variants through substitutions in the antigenic region of the VP1 protein that enable evasion of host population immunity, leading to global spread. In contrast, we show here that the acquisition of new genetic and antigenic characteristics is not the proximal driver of new pandemics. Instead, pandemic GII.4 viruses circulate undetected for years before causing a new pandemic, during which time they diversify and spread over wide geographical areas. Serological data demonstrate that by 2003, some nine years before it emerged as a new pandemic, the ancestral 2012 pandemic strain had already acquired the antigenic characteristics that allowed it to evade prevailing population immunity against the previous 2009 pandemic variant. These results provide strong evidence that viral genetic changes are necessary but not sufficient for GII.4 pandemic spread. Instead, we suggest that it is changes in host population immunity that enable pandemic spread of an antigenically-preadapted GII.4 variant. These results indicate that predicting future GII.4 pandemic variants will require surveillance of currently unsampled reservoir populations. Furthermore, a broadly acting GII.4 vaccine will be critical to prevent future pandemics.

**Significance:** Norovirus pandemics and their associated public health and economic costs could be prevented by effective vaccines. However, vaccine development and distribution will require identification of the sources and drivers of new pandemics. We here use phylogenetics and serological experiments to develop and test a new hypothesis of pandemic norovirus emergence. We find that pandemic noroviruses preadapt, diversify and spread worldwide years prior to emergence, strongly indicating that genetic changes are necessary but not sufficient to drive a new pandemic. We instead suggest that changes in population immunity enable pandemic emergence of a pre-adapted low-level variant. These findings indicate that prediction of new pandemics will require surveillance of under-sampled virus reservoirs and that norovirus vaccines will need to elicit broad immunity.

## Introduction

Noroviruses are the leading cause of acute gastroenteritis in humans worldwide, causing an estimated 684 million gastroenteritis episodes, 200,000 deaths and $65 billion of health and societal costs annually (Kirk et al. 2015; Pires et al. 2015; Bartsch et al. 2016). While more than 30 norovirus genotypes have been described based on sequence variation in the VP1 capsid protein, the GII.4 genotype is responsible for the majority of human cases and outbreaks and has caused six major pandemics since the mid-1990s, each associated with a distinct pandemic variant: US95/96, Farmington Hills 2002, Hunter 2004, Den Haag 2006, New Orleans 2009 and Sydney 2012 (where the year denotes the year of onset of the respective pandemic) (Kroneman et al. 2008; Siebenga et al. 2009; White 2014; Vinje 2015; Motoya et al. 2017; Parra et al. 2017), as well as other geographically-limited outbreaks caused by epidemic variants (Siebenga et al. 2009; Eden et al. 2014).

Vaccines currently under development offer promise to mitigate the global economic and health impact of new GII.4 pandemics (Lindesmith et al. 2015). However, effective vaccine design and distribution depend on understanding the sources from which new pandemic variants emerge and the factors that drive their global circulation especially if, like influenza, vaccine updates are necessary. It has been proposed that new pandemic GII.4 viruses generally evolve from one of the preceding pandemic variants (Siebenga et al. 2007; White 2014) through acquisition of substitutions in the capsid VP1 protein that alter antigenicity and enable evasion of host population immunity (Lindesmith et al. 2008; Lindesmith et al. 2012; Debbink et al. 2013; Lindesmith et al. 2013; Eden et al. 2014). The four most recent pandemic variants caused outbreaks up to five years prior to pandemic emergence (Sdiri-Loulizi et al. 2009; Siebenga et al. 2009; Eden et al. 2014; van Beek et al. 2018), leading to suggestions that both New Orleans 2009 and Sydney 2012 variants circulated at low levels until the acquisition of the VP1 substitutions necessary for pandemic emergence (Eden et al. 2014; White 2014). Recombination has also been proposed to play a role in pandemic emergence (Eden et al. 2013), although precisely how is unknown.

Here, we combine phylogenetic and serological analyses to formulate and test a new hypothesis for GII.4 pandemic emergence. We demonstrate that pandemic GII.4 variants arise years before pandemic spread and diversify over wide geographical areas over the years prior to their emergence. In depth analysis of sequence data shows that genetic substitutions and recombination events that may be important for pandemic emergence are acquired years before such emergence occurs. Serological assays incorporating reconstructed ancestral strains of the Sydney 2012 pandemic variant demonstrate that key antigenic characteristics required for emergence had already been acquired by 2003, nine years prior to pandemic spread. Together, out results show that viral genetic changes (substitutions and/or recombination events) are necessary but not sufficient for pandemic spread and indicate a role for changes in host immunity in the emergence of new variants.

## Results and Discussion

To understand the origins and spread of norovirus pandemics, we reconstructed the temporal history of each genomic region of GII.4. This indicates that GII.4 was present for at least 50 years prior to the first documented pandemic (Figures 1, S1, Table S1), approximately 20 years earlier than previous estimates (Bok et al. 2009) due to inclusion of an additional early sequence that diverges from the root of the phylogeny.

**Figure 1.**
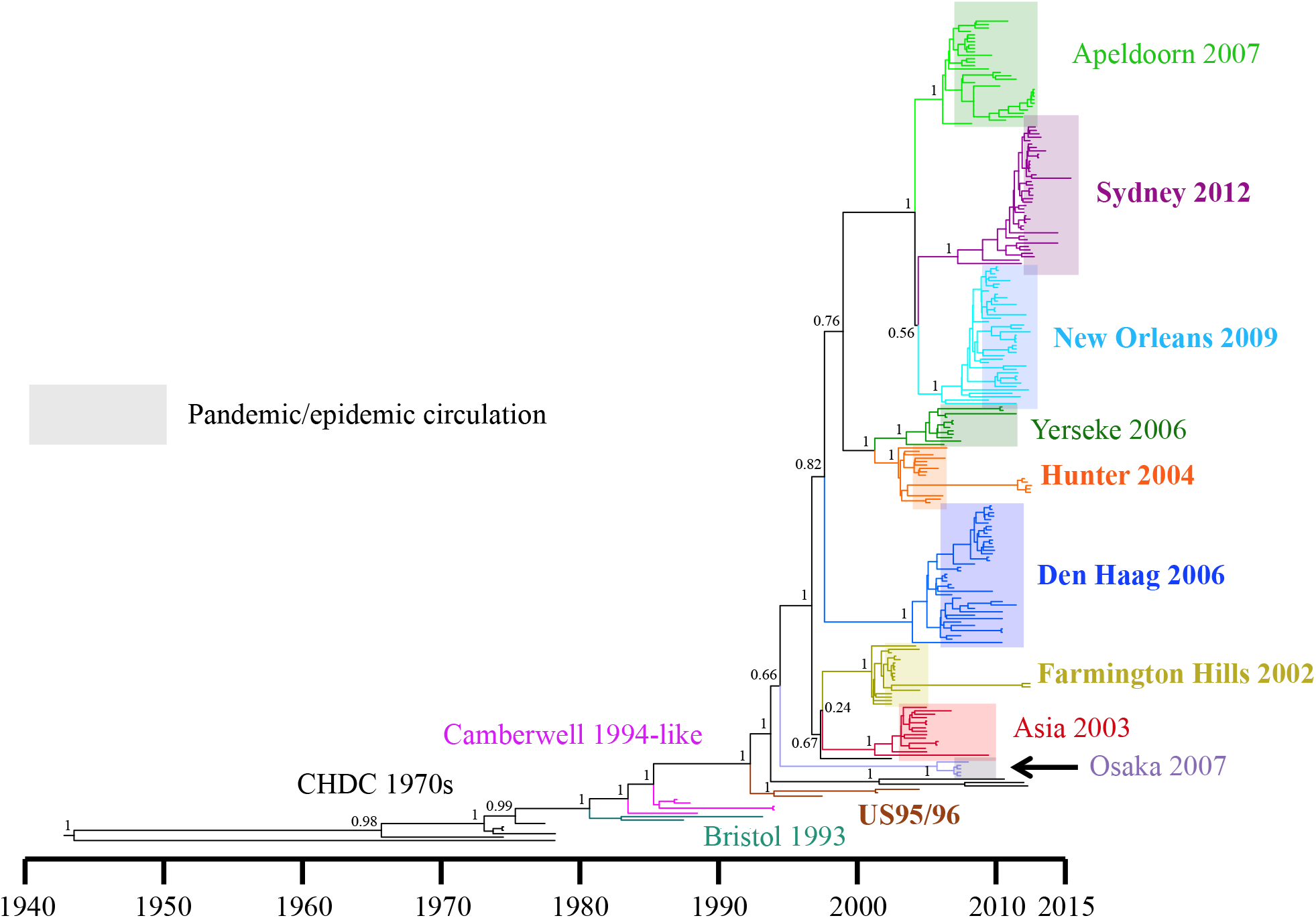
Temporal maximum clade credibility (MCC) tree of GII.4 VP1 sequences from major pandemic and epidemic variants. Variants diverge from all other sampled variants years before their emergence as a pandemic or epidemic (represented by the shaded area). Long branches throughout the tree indicate a high level of unsampled diversity through time. Posterior supports are shown on trunk nodes.

The phylogenetic relationships between GII.4 RdRp, VP1 and VP2 sequences are incompatible with the hypothesis that new pandemic/epidemic viruses evolve from previous pandemic/epidemic variants (Figures 1, S1, Table S2). Instead, the deep phylogenetic nodes suggest that GII.4 variants diverge from one another long before emerging to spread pandemically. The long tree branch lengths indicate ongoing undetected circulation of pre-pandemic variants at low level during the period between divergence and pandemic emergence. For example, Den Haag 2006 and New Orleans 2009 were circulating as unique independent lineages by 1997 (VP1, 95% highest probability density (HPD) 1995-2000) and 2004 (VP1, 95% HPD 2002-2005), respectively. This undetected persistence over long time periods means that multiple future pandemic/epidemic variants co-circulated simultaneously. For example, at least six unsampled lineages co-circulated in the year 2000, four of which gave rise to five subsequent pandemics (Figures 1, Table S2). In support of our findings, we identified and verified previously sequenced pre-pandemic Farmington Hills 2002, Hunter 2004, New Orleans 2009 and Sydney 2012 and pre-epidemic Osaka 2007 sequences (Figure S2, Tables S3, S4). These pre-pandemic sequences sit closer to the root of the clade than sequences collected during the pandemic/epidemic (Figure S2) with placements in the tree in agreement with their sampling dates, supporting our inferred ancestor and divergence dates. Pre-pandemic circulation of Hunter 2004 (Sdiri-Loulizi et al. 2009), Den Haag 2006 (Siebenga et al. 2009; van Beek et al. 2018), New Orleans 2009 and Sydney 2012 (Eden et al. 2014; van Beek et al. 2018) has also been reported. The presence of multiple long future pandemic lineages that are only rarely sampled suggests that highly diverse viral populations are circulating within reservoirs that are not included in current surveillance.

We next used datasets of VP1 sequences to reconstruct a more detailed temporal history of the two most recent pandemic variants, New Orleans 2009 and Sydney 2012 (Figures 2, S3). Similar to other pandemic GII.4 variants (Siebenga et al. 2010), New Orleans 2009 and Sydney 2012 VP1 regions underwent a large increase in relative genetic diversity coinciding with their pandemic emergence in 2009 and 2012, respectively (Figure S3). However, each variant had already diversified into many lineages *prior* to pandemic emergence, at least 67 (95% HPD 41-100) and 88 (95% HPD 59-113) lineages for New Orleans 2009 and Sydney 2012, respectively (Figures 2A-B, S3). Each of the other pandemic GII.4 variants also exhibit this pre-pandemic divergence (Figure 1). Therefore, pandemic variants not only arise long before the pandemic is observed, but also undergo extensive diversification into multiple related lineages that circulate at low levels for years preceding pandemic emergence.

**Figure 2.**
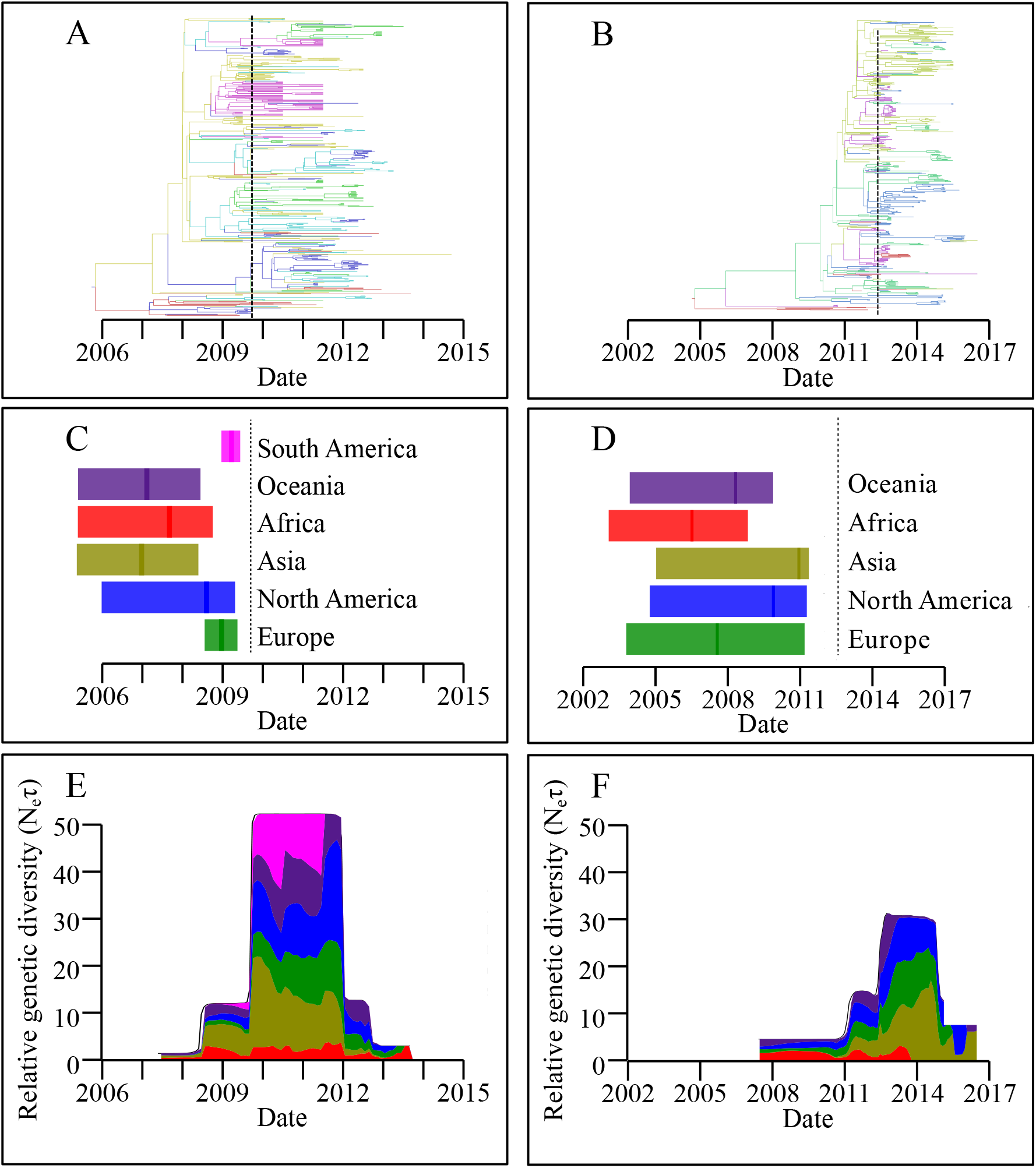
GII.4 variants New Orleans 2009 and Sydney 2012 diversified and spread widely prior to pandemic emergence. (A and B) Spatiotemporally resolved MCC trees for New Orleans 2009 (A) and Sydney 2012 (B) with each branch coloured by inferred location, as in panels C and D. (C and D) Summary of continent import dates for New Orleans 2009 (C) and Sydney 2012 (D); the vertical line is the median import date and the shaded area the 95% HPD. The dashed vertical black line is the inferred date of pandemic emergence. (E and F) Summary of the spatiotemporal distribution of lineages from New Orleans 2009 (E) and Sydney 2012 (F). The proportion of lineages on each continent through time is plotted as a stacked area plot, scaled to the estimated relative genetic diversity.

To determine the extent of circulation prior to pandemic emergence, we reconstructed the spatiotemporal history of New Orleans 2009 and Sydney 2012, which demonstrated entry into each continent prior to pandemic emergence (Figure 2). Specifically, New Orleans 2009 likely entered Africa, Asia and Oceania more than two years prior to pandemic emergence while Sydney 2012 was likely introduced into Africa, Europe and Oceania more than four years before pandemic emergence (Figure 2C-D). Following their introduction, these data suggest sustained intra- and inter-continental circulation of New Orleans 2009 and Sydney 2012 both before (at lower levels) and after (at higher levels) pandemic emergence (Figure 2E-F).

The pandemic variant common ancestor date occurring years prior to pandemic emergence indicates either that the important characteristics for pandemic spread were acquired years before such spread occurred or that such changes were acquired convergently following diversification into multiple lineages. The extent of diversification prior to pandemic emergence argues strongly against the latter scenario. Not only would important changes have to occur in a large number of individual lineages located on multiple continents, but these changes would have to occur approximately simultaneously after a delay of multiple years. We therefore hypothesized that the key characteristics for pandemic spread are acquired by the variant common ancestor years prior to pandemic emergence. To test this hypothesis, we assayed the antigenic properties of two reconstructed Sydney 2012 ancestors: 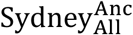, the common ancestor of all sequences genotyped as Sydney 2012 (estimated date: late 2003, 95% HPD early 2000-early 2007) and 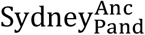, the common ancestor of all pandemic Sydney 2012 viruses (estimated date: late 2008, 95% HPD late 2006-early 2010) (Figure 3A). 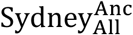 and 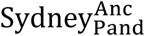 exhibit similar or greater resistance to anti-New Orleans 2009 human polyclonal sera (Figure 3B), mouse polyclonal sera (Figure 3C) and mouse monoclonal antibodies (mAbs, Figure S4) compared with a reference Sydney 2012 virus (Sydney^Ref^) collected during the pandemic. These data indicate that substitutions in VP1 acquired prior to 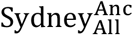 provided resistance to the anti-New Orleans 2009 antibody response at least nine years prior to the onset of the Sydney 2012 pandemic and six years prior to the pandemic emergence of New Orleans 2009.

**Figure 3.**
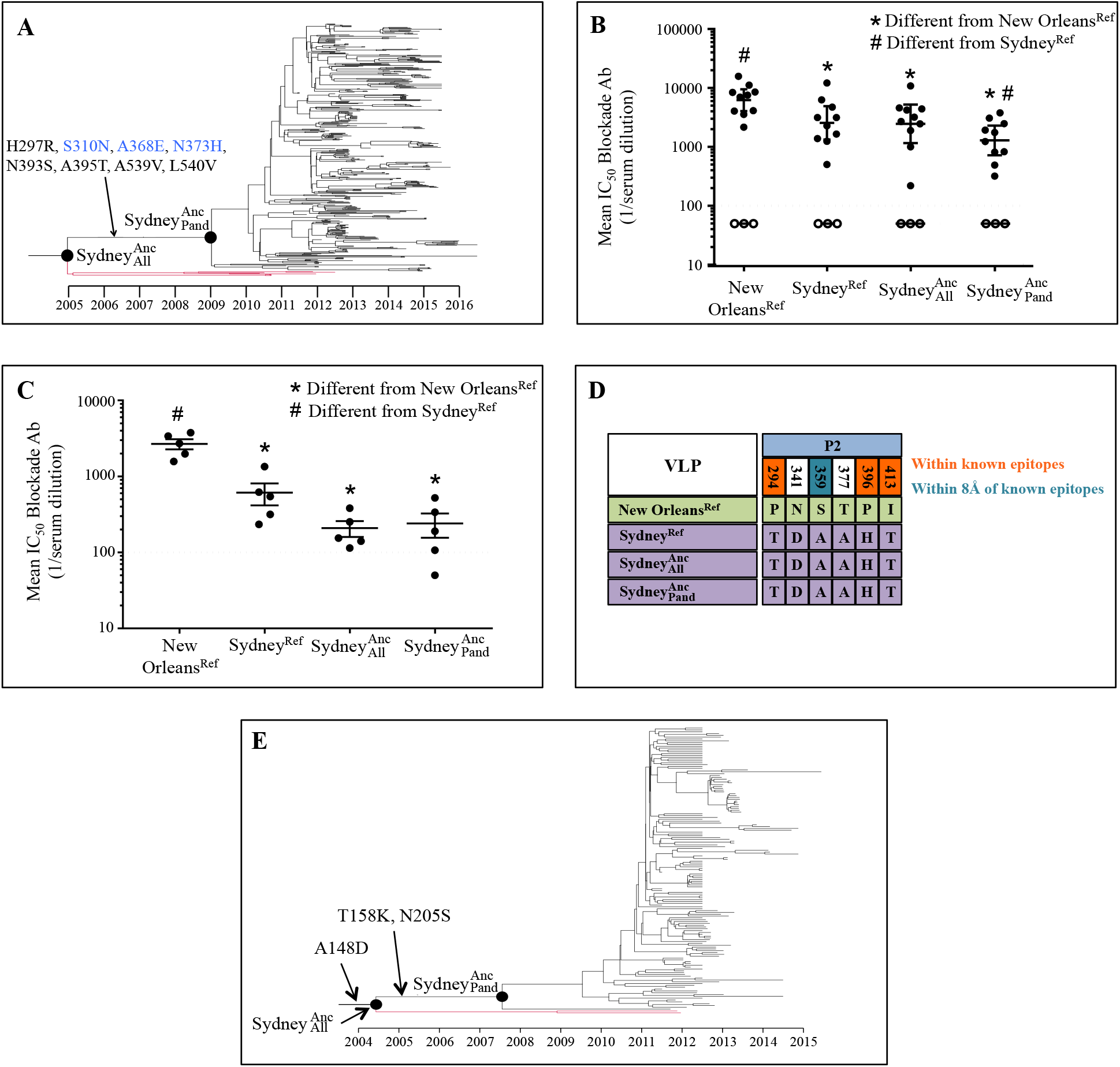
Sydney 2012 could resist anti-New Orleans 2009 immunity by 2003. (A) Temporally resolved Sydney 2012 tree with 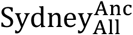 and 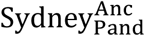 labelled. The lineages that diverged between 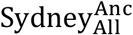 and 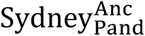 (shown in red) did not persist in the population. Nonsynonymous substitutions that occurred leading to 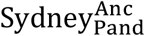 are labelled. Substitutions labelled in blue remained highly conserved in Sydney 2012 (Figure S5). (B) Blockade of 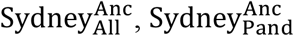, New Orleans^Ref^ and Sydney^Ref^ interaction with pig gastric mucin (PGM) by polyclonal sera from patients infected with New Orleans 2009 (closed circles) or healthy blood donors (open circles, did not block VLPs at the assay limit of detection). Bars are geometric mean values with 95% confidence intervals. Dashed line is assay limit of detection. Statistical significance calculated using the Wilcoxon test. (C) As in B but using polyclonal sera collected from mice exposed to New Orleans 2009. (D) Six amino acid sites in the antigenic VP1 P2 subdomain exhibit a different residue in all three Sydney 2012 VLPs compared with New Orleans 2009. (E) Temporally resolved Sydney 2012 VP2 phylogeny with 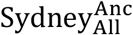 and 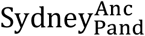 labelled. Nonsynonymous substitutions leading to 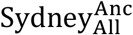 and 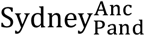 are labelled.

We identified six substitutions in the antigenic P2 domain that could have driven this antigenic change (Figure 3D), including sites within (sites 294, 396 and 413) and structurally close to (site 359) known epitopes (Lindesmith et al. 2012). An additional eight amino acid substitutions occurred in VP1 between 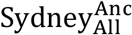 and 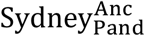 (Figure 3A), of which sites 310 and 368 remain highly conserved within Sydney 2012 (Figure S5), indicating that their acquisition by 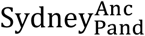 may have been important for its subsequent emergence as a new pandemic. Site 368 was previously demonstrated to alter recognition of mAbs raised against New Orleans 2009 (Debbink et al. 2013) while site 310 is located within the NERK motif that regulates particle breathing and antibody access to epitopes (Lindesmith et al. 2014). The acquisition of these substitutions by 2008 may have been important to enable pandemic emergence years later, by further altering antigenicity or by increasing transmissibility, receptor binding, particle stability or other properties.

We next examined the potential influence of other genomic regions on the pandemic emergence of Sydney 2012. It is unlikely that the nonstructural polyprotein drove Sydney 2012 pandemic spread, as this variant co-circulated with the nonstructural polyprotein from the unrelated GII.Pe, GII.P4 and (more recently) GII.P16 genotypes (Figure S6) (Wong et al. 2013; Ruis et al. 2017). Substitutions occurred within VP2 leading to 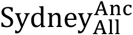 (site 148) and 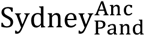 (sites 158, 205) and remained highly conserved within the Sydney 2012 clade (Figures 3E, S5). Without better structural or functional characterization of the VP2 protein, interpreting the contribution of these changes is difficult. However, the 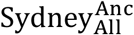 and 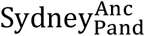 VP2 proteins occurred in mid 2004 (95% HPD mid 1999-early 2009) and early 2008 (95% HPD mid 2005-early 2010), respectively (Figure 3E). Therefore, as with VP1, if these substitutions were important for pandemic emergence, they were acquired years earlier.

A similar process occurred for the other four pandemic GII.4 variants to have emerged since 2002, where in each case, key nonsynonymous substitutions in the nonstructural polyprotein, VP1 and VP2 that characterize the new pandemic variant occurred along the branch leading to the respective common ancestor long before these variants spread pandemically (Tables S5, S6). The fact that the VP1 substitutions included sites within known blockade epitopes (Lindesmith et al. 2012) provides further support for the hypothesis that the antigenic changes required for pandemic emergence were acquired years before pandemic emergence actually occurred.

While recombination has previously been suggested to drive GII.4 pandemics (Eden et al. 2013), we find that each recombination event in which a variant acquired a new nonstructural polyprotein or VP2 occurred years prior to pandemic emergence (Table S7, Supplementary text). These results indicate that recombination events are not the proximate drivers of new norovirus pandemics.

Together, our results indicate that pandemic GII.4 variants arise, diversify and spread widely years before they emerge to cause a pandemic. If a new pandemic was triggered by antigenic change or another viral characteristic, a single lineage would rapidly increase in prevalence, as is observed for influenza A H3N2 (Bedford et al. 2015). Our data instead strongly support a scenario where the key antigenic and other changes are acquired through substitutions and/or recombination events years before pandemic emergence. Therefore the pandemic event is not proximally driven by genetic changes in any of the genomic regions altering antigenicity, receptor binding or another property.

These results raise the question of what drives a variant that has been circulating widely and cryptically for years to suddenly increase in frequency, dominate outbreaks worldwide and rapidly replace the preceding pandemic variant. It is unlikely that stochastic factors would enable widely spread lineages from a single variant within an individual genotype to emerge simultaneously. We therefore suggest that norovirus pandemics are driven by a change in host factors. Given the importance of herd immunity in variant emergence (Lindesmith et al. 2012; Debbink et al. 2013; Lindesmith et al. 2013), we hypothesize that a shift in host population immunity (potentially driven by growing immunity against the previous pandemic variant) opens a population-wide immunological niche into which the multiple circulating, but hidden, lineages of the new pandemic variant can expand, having acquired the necessary antigenic characteristics to do so years before. Therefore viral genetic changes are necessary but not sufficient for pandemic emergence. Instead, a shift in host immunity combines with antigenic preadaptation to drive a new pandemic. Vaccination against norovirus has previously been demonstrated to alter the serological blockade repertoire (Lindesmith et al. 2015), supporting the notion that infection of a large number of individuals can alter population-level immunity. Alternatively, variant emergence could be delayed by widespread heterotypic immunity that decays faster than homotypic immunity raised against the preceding pandemic variant. A natural corollary is that future pandemic GII.4 variants are continuing to circulate and diversify undetected within the reservoir until such time as changes in host immunity should favor the emergence of a new pandemic variant.

These results also raise the question of where pre-pandemic variants circulate over the years prior to emergence. Both immunocompromised patients and animals have been mooted as a potential source of pandemic GII.4 variants (Kari Debbink et al. 2014; Karst and Baric 2015). While it is possible that either of these hosts could be the source of the ancestral variant, both seem unlikely to be the source of the diversifying pre-pandemic lineage. It is unlikely that multiple pandemic lineages could emerge simultaneously from a single immunocompromised host at pandemic onset. It is also difficult to explain how multiple immunocompromised patients could form the inter-connected intercontinental transmission network required for this pre-diversification, supporting recent suggestions that immunocompromised patients are an unlikely reservoir (Eden et al. 2017). Additionally, while GII.4 viruses have occasionally been detected in stool samples from cows, pigs and dogs (Mattison et al. 2007; Summa et al. 2012), concurrent emergence of multiple lineages would require multiple zoonotic transmissions and no such transmissions have been observed (Wilhelm et al. 2015). A more parsimonious explanation for our findings is that pandemic GII.4 variants circulate within the community and are not detected by current surveillance efforts that largely target outbreaks, predominantly in hospital and institutional settings (Inns et al. 2017). More extensive co-circulation of viral lineages has been noted in influenza A H1N1 and influenza B compared with influenza A H3N2 and has been correlated with slower rates of antigenic drift and a lower average age of infection (Bedford et al. 2015; Vijaykrishna et al. 2015). While noroviruses are prevalent in individuals of all age groups, the infection rate is highest in young children (Lopman et al. 2016; O’Brien et al. 2016). In this regard, of the 16 identified pre-pandemic samples for which this information is available, 15 are either from children or were sampled in a nursery or primary school (Figure S2, Table S3). The ability of the ancestral Sydney 2012 VLPs to evade anti-New Orleans 2009 polyclonal sera raised in mice that have not previously been exposed to norovirus (Figure 3C) suggests that these viruses would have been able to evade immunity raised in young children. Thus, while continued strain monitoring, as captured by norovirus outbreak surveillance networks such as CaliciNet (Vega et al. 2011) and NoroNet (van Beek et al. 2018), has significant value for vaccine development, additional efforts should focus on identifying potential reservoirs from which future pandemic norovirus variants could emerge, including symptomatic and asymptomatic children (Rouhani et al. 2016) in healthcare and community settings. The antigenic preadaptation of future pandemic variants suggests that such a surveillance system combined with antigenic testing could efficiently predict currently low-level variants that might become pandemic in the future. It will also be key to understand how these variants interact with prevailing population immunity.

Our results also have important implications for current efforts to develop norovirus vaccines. Should the new pandemic variant emerge from the preceding variant, a vaccine targeting the current variant may prevent the next pandemic. However, under our hypothesis, a vaccine that boosts immunity to the current variant may actually hasten emergence of the next pandemic. It is therefore essential that norovirus vaccines provide broad immunity against GII.4 viruses (Lindesmith et al. 2015).

## Materials and Methods

### Reconstruction of the temporal history of GII.4 norovirus

We collected all norovirus sequences available on GenBank as of 30/10/2015 containing the complete RNA-dependent RNA polymerase (RdRp), VP1 and VP2 genome regions. Each sequence was genotyped using the norovirus genotyping tool (Kroneman et al. 2011) and the 871 sequences containing the GII.4 VP1 were retained. Due to inter-genotype recombination events close to the ORF1-ORF2 boundary (Eden et al. 2013), the dataset contains sequences with the GII.P1, GII.P4, GII.P12 and GII.Pe RdRps.

Due to the presence of recombination close to ORF boundaries, we ran all analyses on the RdRp, VP1 and VP2 separately. Each genome region was aligned at the amino acid level using MUSCLE (Edgar 2004). We screened for the presence of intra-genic recombination in the RdRp, VP1 and VP2 separately using the Single Breakpoint (SBP) method implemented in HyPhy (Pond et al. 2006), identifying 19 sequences as potential recombinants in VP1 (Table S8). These sequences clustered differently with strong support on either side of the putative breakpoint in phylogenetic trees reconstructed with RAxML v8.1 (Stamatakis 2014) with the GTR model and gamma rate heterogeneity with four gamma classes. SBP was then run on the alignment again following removal of these samples to ensure the recombination signal had been removed. A summary of the number of remaining sequences from each GII.4 variant and each RdRp genotype is shown in Table S9. The variant names used here are those returned by the norovirus genotyping tool (Kroneman et al. 2011).

Methods employing sequence data and sampling dates to infer divergence times are valid only if there is a temporal evolutionary signal in the dataset (Rambaut et al. 2016). To assess whether each of our datasets exhibits a temporal evolutionary signal, we reconstructed a maximum likelihood tree using RAxML, as above. We identified the best-fitting root position using TempEst v1.5 (Rambaut et al. 2016) and calculated the R2 correlation between root-to-tip distance and sampling date (Figure S7). There was a significant temporal evolutionary signal within each dataset (p < 0.001).

We reconstructed the temporal evolutionary history of each genomic region using the Bayesian Markov chain Monte Carlo approach implemented in BEAST version 2.2.1 (Bouckaert et al. 2014). Analyses were run independently on the RdRp, VP1 and VP2. As the Den Haag 2006 and New Orleans 2009 variants have a greater number of sequences compared with other variants (Table S9), we took three random subsamples of 41 sequences from each of these variants (with 41 chosen to match the number of sequences in the third most numerous variant); results were insensitive to the subsampled dataset (Table S10). Each sequence was labelled with the most accurate collection date possible: the day of collection if available, the month of collection if the day of collection was not available or the year of collection if the month of collection was not available. Each dataset was analyzed using the GTR substitution model with gamma rate heterogeneity and partitioned so codon positions 1 and 2 shared a substitution model and codon position 3 had a different substitution model. We employed both the strict and relaxed lognormal clock models to examine variation in the substitution rate within each dataset. We used a lognormal prior distribution for each dataset with mean 4.3×10^−3^ substitutions/site/year and standard deviation 0.1 for the VP1 (Bok et al. 2009) and with mean 4.32×10^−3^ substitutions/site/year and standard deviation 0.1 for the RdRp (Siebenga et al. 2010). At the time of our analysis, there was no previously published substitution rate for VP2 across the GII.4 clade. We therefore employed the same prior as for the VP1 dataset. We applied a coalescent Bayesian skyline tree prior. Three replicate runs with different starting values were performed for each dataset and clock model and run until convergence, as assessed using Tracer v1.6 (Rambaut et al. 2014). The replicate runs were combined with removal of suitable burnin using LogCombiner v2.2.1 and maximum clade credibility trees were obtained using TreeAnnotator v2.2.1. In each case there was strong support to reject the strict clock model in favor of the relaxed lognormal clock model. Therefore the results employing the relaxed lognormal clock model were used for all further analyses.

We estimated variant divergence dates and recombination dates by combining the posterior distribution of trees from each subsampled dataset into a single posterior distribution. We inferred variant divergence dates by calculating the date of the most recent common ancestor between each pair of variants in each tree in this posterior distribution. To calculate the date of each recombination event, we identified the mean and 95% HPD of the distribution of branch start and end times of the corresponding branch in each tree in the posterior distribution.

### Identification of pre-pandemic and pre-epidemic GII.4 sequences

We defined pre-pandemic/pre-epidemic sequences as those that cluster with a GII.4 variant but were collected prior to the year in which that variant emerged pandemically or epidemically. We genotyped all GII.4 VP1 sequences present on GenBank containing more than 400 nucleotides (Kroneman et al. 2011). We identified 50 sequences with a reported collection date earlier than the year of pandemic/epidemic emergence of the respective variant. We estimated the collection date of each of these sequences using BEAST v2.4.2 (Bouckaert et al. 2014), assuming a uniform prior distribution with minimum 1974.5 and maximum 2015.446575 (the dates of the earliest and latest collected sequences in the main dataset). The 95% HPD of the estimated collection date overlapping with the reported collection date was taken as evidence to support, but not confirm, the reported collection date.

### Reconstruction of the temporal history of New Orleans 2009 and Sydney 2012

We compiled expanded datasets from GenBank containing 460 and 533 P2 domain sequences for New Orleans 2009 and Sydney 2012, respectively. Each dataset was aligned and inferred to exhibit a temporal evolutionary signal (p < 0.001) as above. We reconstructed the evolutionary dynamics of New Orleans 2009 and Sydney 2012 independently using BEAST v2.2.1 (Bouckaert et al. 2014), using the HKY nucleotide substitution model with four gamma classes. We employed a lognormal prior on the substitution rate with mean 6.83×10^−3^ and standard deviation 0.1 to accommodate the mean and 95% HPD of our estimate of the substitution rate across the complete GII.4 VP1 clade. We applied the strict and relaxed lognormal clock models and found strong support (log10 Bayes factor > 100) to reject the strict clock model in favor of the relaxed lognormal clock model in each case. We therefore used the results from the relaxed lognormal clock runs in all further analyses. Bayesian skyline plots were reconstructed using Tracer v1.5. The date at which the Bayesian skyline plot exhibits a large increase in relative genetic diversity was used as the time of pandemic onset. The number of lineages present at the onset of the pandemic was calculated as the number of lineages present at this point in time in each tree in the posterior distribution.

### Reconstruction of the spatiotemporal history of New Orleans 2009 and Sydney 2012

We collected datasets containing all available VP1 sequences on GenBank as of 09/02/2017 from New Orleans 2009 (n=565) and Sydney 2012 (n=708). Each dataset exhibited temporal signal as above. Examination of the sampling locations showed that there was typically only a small number of sequences from each country (Table S11). We therefore used the continent of collection as the location label. The New Orleans 2009 dataset contained a large number of sequences from Asia and Oceania relative to the other continents, while the Sydney 2012 dataset contained a large number of sequences from Asia relative to the other continents (Table S11). Should sequences from the same continent cluster together within the tree, an excess of sequences from one continent is unlikely to alter estimates of ancestral locations. However, if sequences from the different continents are typically interspersed within the tree, an excess of sequences from one continent could result in artifactual support for this continent being the location of ancestral nodes. The sequences from each continent are interspersed throughout the tree in both New Orleans 2009 and Sydney 2012 (Figure S8). We therefore randomly down-sampled New Orleans 2009 sequences from Asia and Oceania and Sydney 2012 sequences from Asia to match the number of sequences from the next most commonly represented continent. We carried out three random subsamples and performed all analyses on each subsampled dataset. All results were insensitive to the subsampled dataset.

We used discrete phylogeography (Lemey,Philippe et al. 2009) implemented in BEAST v2.4.2 (Bouckaert et al. 2014) to reconstruct the spatiotemporal history of New Orleans 2009 and Sydney 2012. Sequences were labelled with the most accurate collection date possible. We modelled the nucleotide substitution process using the HKY substitution model and gamma rate heterogeneity with four gamma classes. We applied both the strict and relaxed lognormal clock models. In each case there was strong support to reject the strict clock model in favor of the relaxed lognormal clock model (log10 Bayes factor 66-83) and we therefore used the relaxed lognormal clock model for our inferences. However, the results with the strict clock model are qualitatively very similar, indicating that potential over-parameterization due to the large number of branch-specific rates with the relaxed lognormal clock model has not influenced our results. We applied a lognormal prior on the substitution rate with mean 7×10^−3^ for New Orleans 2009 and 6.4×10^−3^ for Sydney 2012 and standard deviation 0.1 in each case, based on the posterior estimates of the variant substitution rates inferred previously. We employed a Bayesian coalescent skyline tree prior for each dataset. We applied a discrete phylogeographic model to describe lineage migrations within each dataset *(24)*, consisting of a symmetric transition matrix for the migration rates and a set of Bayesian stochastic search variable selection (BSSVS) indicator variables. We assumed a Poisson prior for the number of ‘on’ BSSVS variables with mean 5 for the New Orleans 2009 datasets and mean 4 for the Sydney 2012 datasets. We used an exponential prior with mean 1.0 migration rate per lineage per year for the overall rate of geographical transition and a gamma prior with shape 1.0 and scale 1.0 for each of the relative geographical transition rates. Three replicate runs with different starting parameters were carried out with each dataset and run until convergence, as assessed using Tracer v1.6 (Rambaut et al. 2014). Runs from each subsampled dataset were combined into a single posterior distribution for downstream analyses.

We identified the date of first import into each continent by calculating either the root date if the continent was inferred to be the root location or the earliest branch midpoint where the downstream node was inferred to be within the continent. We calculated this date within each tree in the posterior distribution. We therefore assume that migration events occurred at the midpoint of the branch. We obtain similar results using the earliest non-root node inferred to be within the continent, which assumes that migration occurs at the end of the branch. We used the program posterior analysis of coalescent trees (PACT) v0.9.4 to compute the proportion of lineages present on each continent through time.

### Reconstruction of ancestral Sydney 2012 viruses and identification of substitutions leading to each GII.4 variant

We collected a dataset containing all 2198 available GII.4 VP1 sequences, including sequences from all of the major GII.4 variants (Table S12). Alignment and phylogenetic reconstruction was carried out as above. We used RAxML to optimize branch lengths within the maximum likelihood phylogenetic tree and ten bootstrap tree topologies using the amino acid alignment and the WAG substitution model with optimized base frequencies. We used multiple tree topologies to assess the influence of tree topology on our inferences. We carried out ancestral reconstruction at the amino acid level with PAML v4.9 (Yang 2007) using the WAG substitution model and optimized base frequencies. The ancestral sequence at 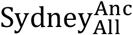 and 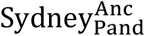 was identical with each tree topology and the residue inferred at each site was supported by posterior probability > 0.95. We identified VP1 substitutions by comparing the sequence of the variant root ancestor with the sequence of the immediately upstream node in each tree.

To identify substitutions with the nonstructural polyprotein and VP2 we collected datasets containing all available GII and GIV nonstructural polyprotein sequences (Table S13) and all available GII.4 VP2 sequences (Table S12), respectively. We identified substitutions leading to each GII.4 variant using the same process described for VP1 above.

### Surrogate neutralization assay (Antibody blockade assay)

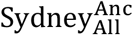 and 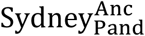 VP1 genes were codon optimized for mammalian expression and synthesized by Bio Basic Inc. (Amherst, NY) and VLPs were expressed in baby hamster kidney cells from Venezuelan equine encephalitis virus replicons *(11)*. 0.25 μg/ml VLP was pretreated with decreasing concentrations of antibody/serum for 1 h before addition to pig gastric mucin type III (PGM, Sigma Aldrich, St. Louis, MO) coated plates for 1 h. Bound VLP was detected with anti-GII.4 2012 ref rabbit hyperimmune sera. Percent control binding is defined as the binding with antibody/serum pretreatment compared to the binding without multiplied by 100. All incubations were done at room temperature. The blockade data were fit using sigmoidal dose-response curve analysis of nonlinear data in GraphPad Prism 702 and IC50 titers with 95% confidence intervals calculated. Antibodies that did not block 50% of binding at the highest dilution tested were assigned an IC50 of 2 times the assay limit of detection for statistical comparison *(11)*. Anti-New Orleans 2009 polyclonal sera was collected from ten patients naturally infected during a New Orleans 2009 outbreak in a long-term care facility in March 2010. Infection was confirmed by symptoms and norovirus detection in acute stool. The human serum samples were collected from a study that was approved by institutional review board of the Centers for Disease Control and Prevention (CDC Protocol #5051). Mouse anti-GII.4 New Orleans 2009^Ref^ sera were generated in immunized mice as described in (K. Debbink et al. 2014). Mouse monoclonal antibodies to GII.4 New Orleans 2009^Ref^ (Lindesmith et al. 2013) were generated by VLP immunization.

### Data availability

All alignments, phylogenetic trees and BEAST XML files will be made available on GitHub on acceptance of the manuscript. VLPs are available from the authors.

## Acknowledgments

The authors would like to thank Victoria Madden of Microscopy Services Laboratory, Department of Pathology and Laboratory Medicine, University of North Carolina-Chapel Hill for expert technical support. The authors are grateful to Oliver G. Pybus of Department of Zoology, University of Oxford for critical review of the manuscript. This work was supported by the Wellcome Trust (grant number 203268/Z/16/Z to J.B), a studentship from UCL CoMPLEX (to C.R), the National Institute for Health Research UCL-UCLH Biomedical Resource Centre, the Medical Research Council (grant number U117573805 to R.A.G), the National Institutes of Health, Allergy and Infectious Diseases (grant number U19 AI109761 to R.S.B), the EU FP7 PATHSEEK grant and a Medical Research Council studentship. The funders had no role in study design, data collection and interpretation, or the decision to submit the work for publication. The findings and conclusions in this article are those of the authors and do not necessarily represent the official position of the Centers for Disease Control and Prevention.

