## Supplemental figures, tables and text for "Preadaptation of pandemic GII.4 noroviruses in hidden virus reservoirs years before emergence"

### Supplementary Text, Figures and Tables

#### Supplementary Text

*Intergenic recombination*

Recombination between noroviruses frequently occurs close to open reading frame boundaries , including within GII.4 (Eden et al. 2013). We observe extensive topological differences between the RdRp, VP1 and VP2 trees in well supported regions, concurring with this previous evidence. In addition to the previously reported acquisition of a Den Haag 2006-like VP2 by the Osaka 2007 VP1 (Eden et al. 2013), we also find that that Apeldoorn lineage acquired a Yerseke 2006-like VP2 in 2003, prior to diverging into the Apeldoorn 2007, New Orleans 2009 and Sydney 2012 variants (Figure S9, Table S7). Multiple recombination events have occurred between the RdRp and VP1. Consistent with previous results (Eden et al. 2013), we infer that Asia 2003 acquired a GII.P12 RdRp and Osaka 2007 acquired a GII.Pe RdRp (Table S7). Multiple recombination events are required to explain the acquisition of RdRp and VP2 regions by the Apeldoorn lineage VP1 (Figure S9). A common ancestor of the New Orleans 2009 and Sydney 2012 (and possibly the Apeldoorn 2007) variants acquired a Yerseke 2006-like RdRp in 2004 (Figure S9, Table S7). While it was not possible to conclusively resolve the recombination events within the Apeldoorn lineage, Figure S9 depicts two plausible scenarios. As previously reported, the Sydney 2012 variant circulated commonly with both the GII.P4 New Orleans 2009-like RdRp and the GII.Pe RdRp (Wong et al. 2013). At least three independent recombination events are required to explain the distribution of sequences with the GII.Pe RdRp in the Sydney 2012 VP1 tree (Figure S6). Importantly, each of the recombination events where the two contributing variants could be identified occurred years prior to the pandemic or epidemic emergence of the contributing variants (Table S7).

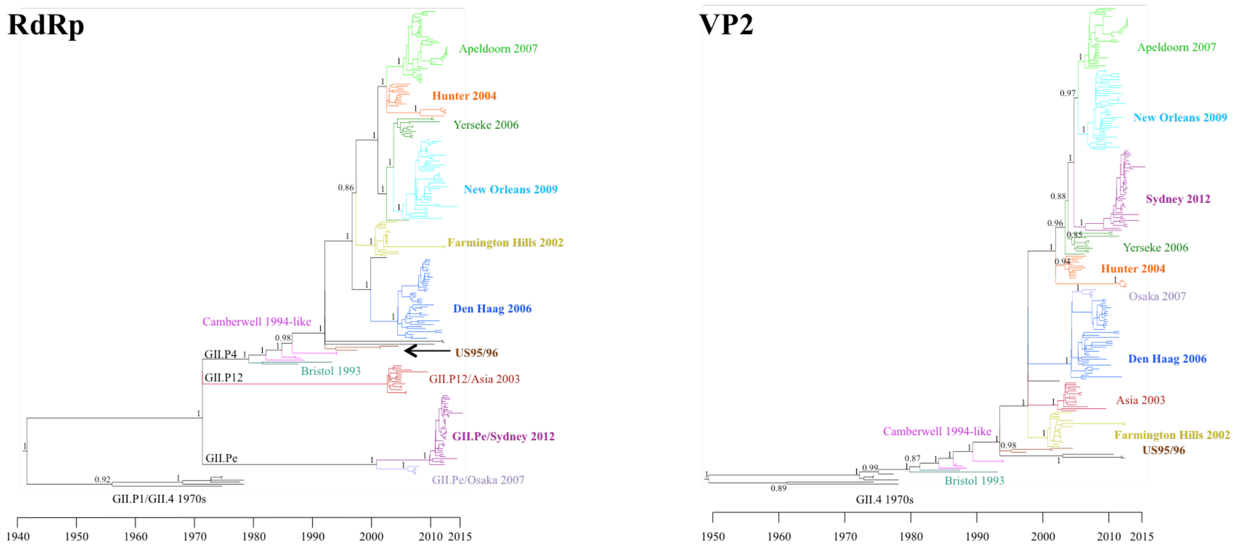


**Fig. S1.**

Temporal MCC trees of the RdRp and VP2. The temporal history of the RdRp and VP2 was reconstructed using BEAST. As with the VP1 tree in Figure 1, the trees of these genomic regions exhibit a high degree of unsampled diversity with a large number of long branches. Each variant diverged from all other sampled variants years prior to pandemic/epidemic emergence. Variants are labelled in different colors. Posterior supports are shown on trunk nodes.

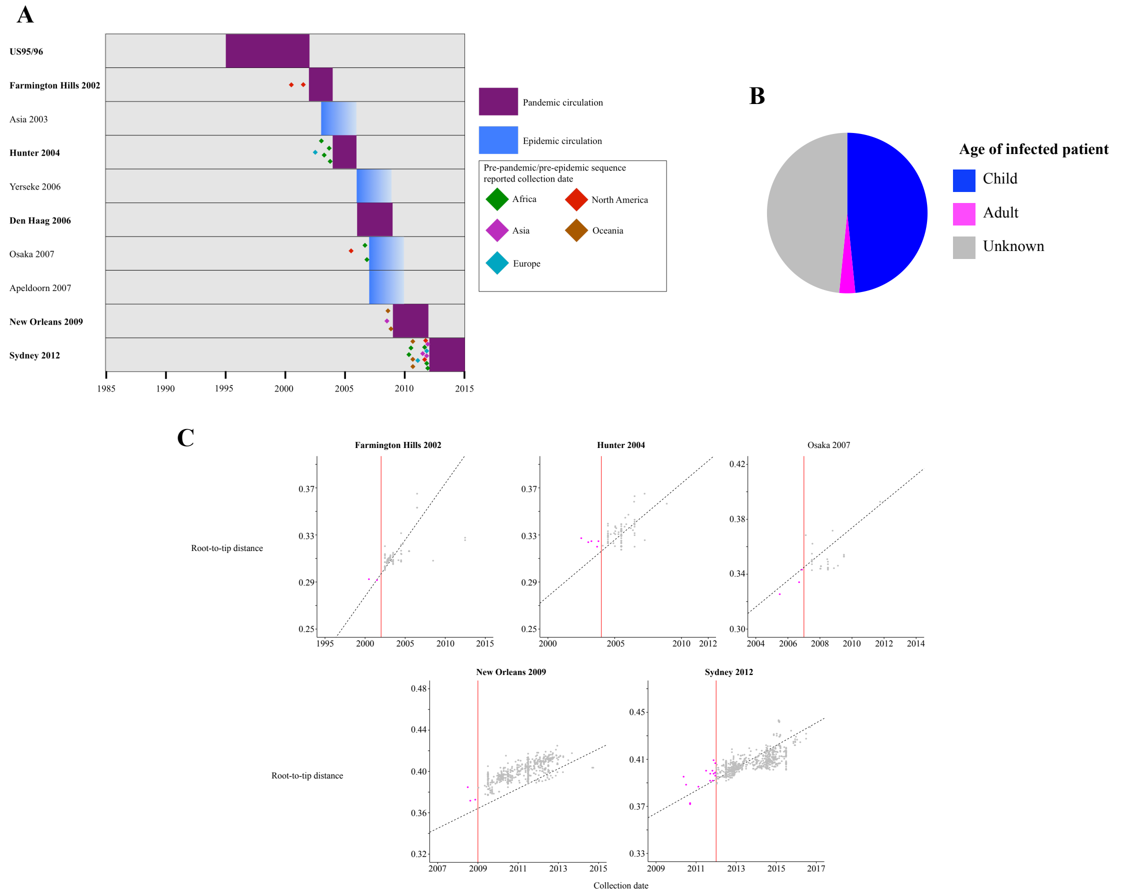


#### Fig S2

(**A**) Identification of pre-pandemic and pre-epidemic sequences. We identified all available GII.4 norovirus sequences with a reported collection date earlier than the start of the year of pandemic/epidemic emergence of the genotyped variant. We verified the collection date of each sequence using Bayesian tip dating (see methods) and identified 31 pre-pandemic/pre-epidemic sequences from the Farmington Hills 2002, Hunter 2004, Osaka 2007, New Orleans 2009 and Sydney 2012 variants. Each sequence is represented here by a diamond at the date at which the sequence was collected, with the color of the diamond representing the continent on which the sequence was collected. The shaded area represents the period of pandemic (purple) or epidemic (blue) circulation. (**B**) The age of the infected patient was reported for 16 of the 31 pre-pandemic/pre-epidemic sequences. 15 of these 16 sequences were collected from children. (**C**) Putative pre-pandemic/pre-epidemic sequences exhibit a level of divergence consistent with their reported collection date. We reconstructed a nucleotide maximum likelihood tree on all available GII.4 VP1 sequences and rooted to maximize the correlation between root-to-tip distance and collection date. Each panel contains all of the sequences from the corresponding variant. Putative pre-pandemic/pre-epidemic sequences are shown in magenta, the remaining sequences are shown in grey. The black dashed line is a regression line between root-to-tip distance and collection date calculated on all GII.4 VP1 sequences. The red vertical line represents the start of the year in which the variant emerged as a pandemic or epidemic.

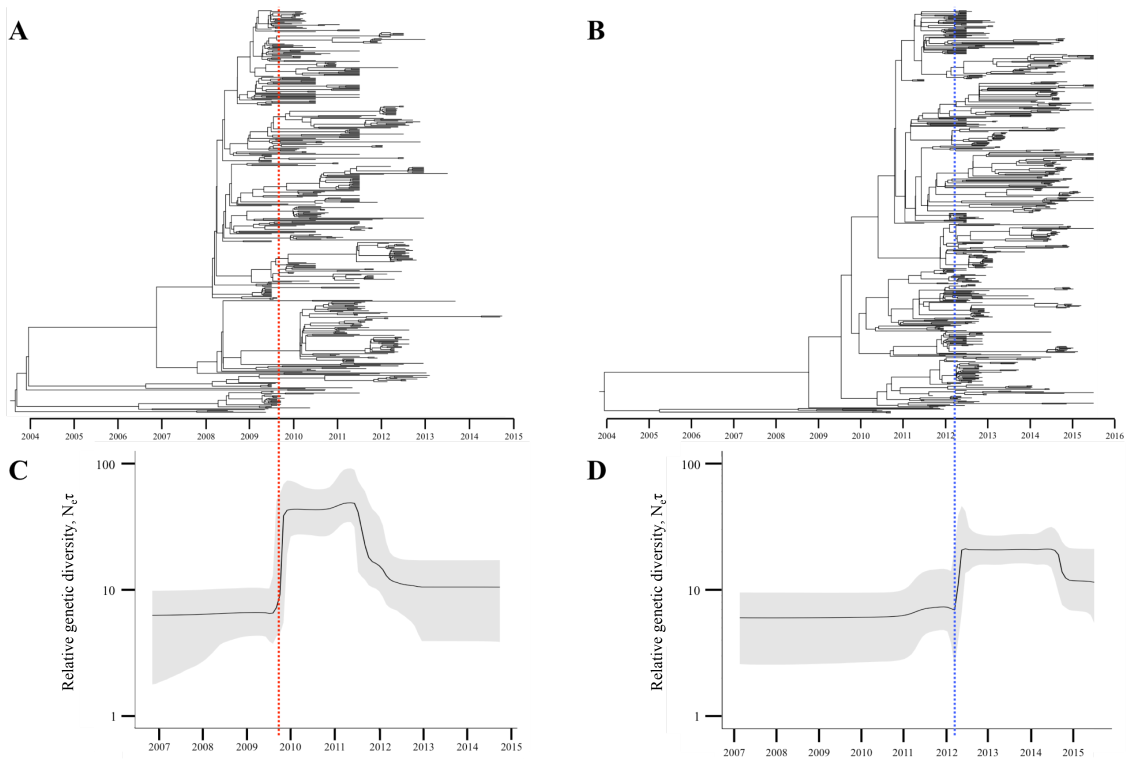


**Fig. S3**
Evolutionary dynamics of New Orleans 2009 and Sydney 2012. Summary of phylodynamic analyses of New Orleans 2009 (A and C) and Sydney 2012 (B and D). (A and B) Temporal MCC trees of all available New Orleans 2009 (n=466) and Sydney 2012 (n=533) P2 domain sequences reconstructed using BEAST. (C and D) Bayesian skyline plots showing a measure of relative genetic diversity through time. The solid black line is the median value and the grey shaded area the 95% HPD. The vertical red and blue lines represent the time of onset of the New Orleans 2009 and Sydney 2012 pandemics, as estimated from the Bayesian skyline plots. By this time each variant had already diverged into a large number of lineages.

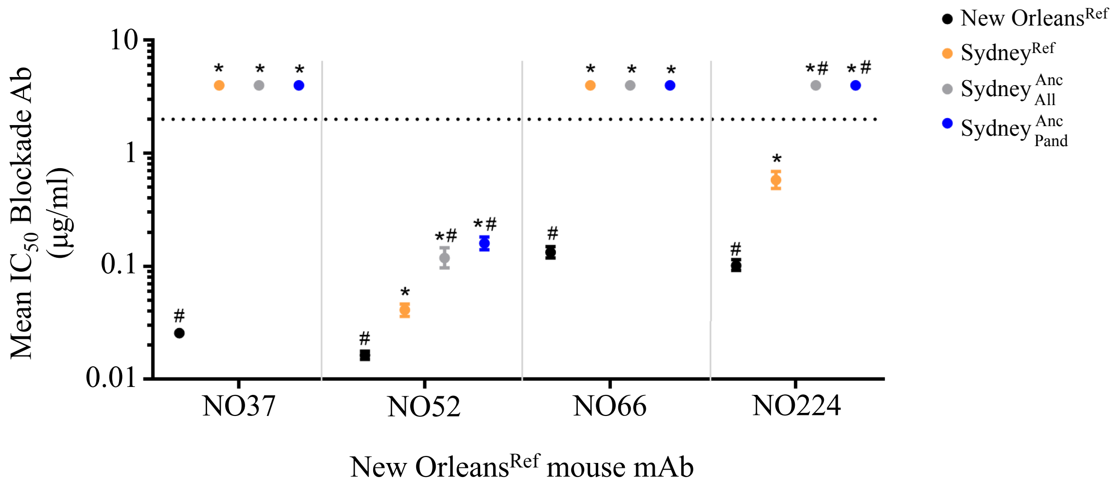


**Fig. S4.**

Sydney 2012 could resist anti-New Orleans 2009 murine mAbs by 2003. We tested the ability of mouse mAbs raised against New Orleans 2009 to block interaction of $Sydney^{Anc}_{All}$, $Sydney^{Anc}_{Pand}$, New $Orleans^{Ref}$ and $Sydney^{Ref}$ VLPs with pig gastric mucin (PGM). The ancestral Sydney 2012 VLPs resisted mAbs raised against blockade epitopes of New $Orleans^{Ref}$ to a comparable or greater degree compared with the $Sydney^{Ref}$ VLP. Markers represent the mean and error bars the 95% confidence intervals. * significantly different from New $Orleans^{Ref}$, # significantly different from $Sydney^{Ref}$ (Dunnett multiple comparison test).

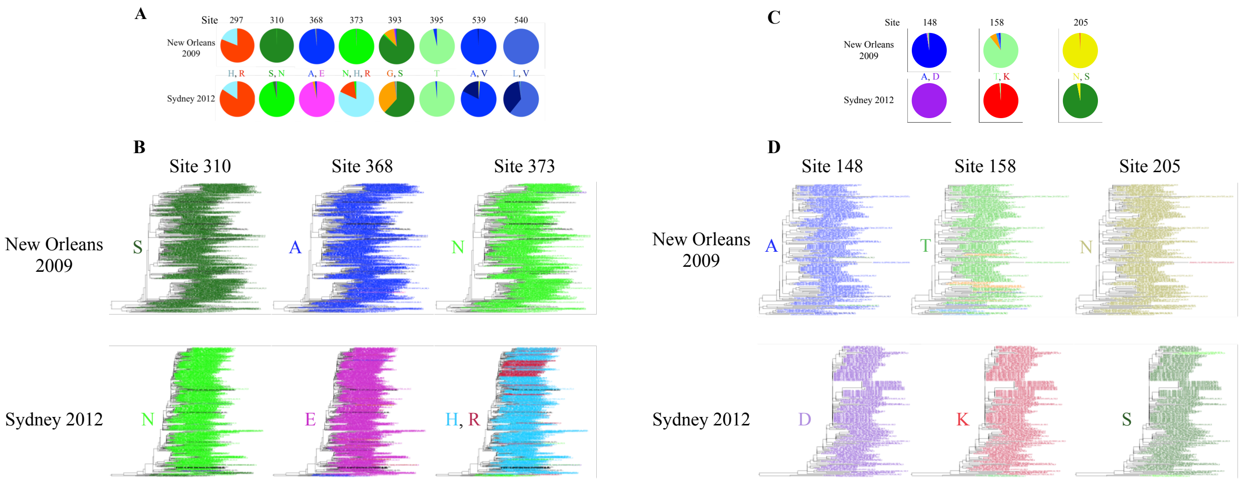


**Fig. S5.**

Substitutions potentially important for the pre-adaptation of Sydney 2012. (A and C) We identified the distribution of amino acid residues within New Orleans 2009 and Sydney 2012 at each site in VP1 (A) and VP2 (C) that underwent a substitution leading to Sydney 2012. The residues at each site are shown between the respective pie charts. In VP1, only sites 310, 368 and 373 exhibit different amino acid residues in Sydney 2012 compared with New Orleans 2009. (B and D) Conservation of sites across the Sydney 2012 clade. Each tip within maximum likelihood trees of all available New Orleans 2009 and Sydney 2012 VP1 sequences is colored by the amino acid residue within that sequence at the respective site. Each of the sites is conserved downstream of Sydney$^{\mathrm{Anc}}_{\mathrm{Pand}}$, with the exception of VP1 site 373 which changes between histidine and arginine, likely on multiple occasions. Black tips indicate that the site was not sequenced within that sample.

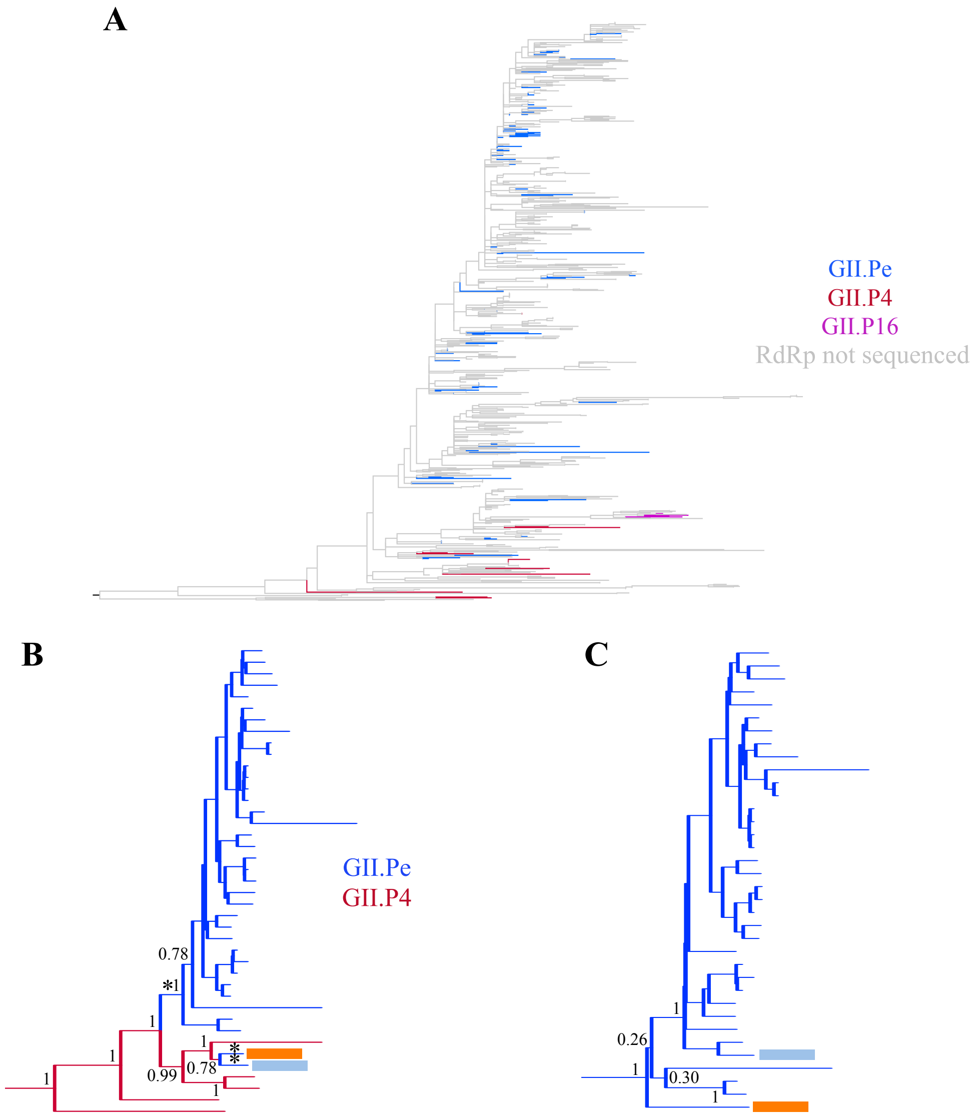


**Fig. S6.**
(A) The tips in the Sydney 2012 VP1 tree are coloured by the associated nonstructural polyprotein genotype. This variant co-circulated with the GII.Pe and GII.P4 nonstructural polyproteins throughout its pandemic period and has more recently circulated with the GII.P16 nonstructural polyprotein. (B) The Sydney 2012 clade in the VP1 tree in Figure 1 is shown. Tip branches are colored by the RdRp found with that sequence: red – GII.P4 New Orleans 2009-like, blue – GII.Pe. Acquisition of the GII.Pe RdRp was inferred to have occurred along the three branches marked with asterisks. (C) The GII.Pe-Sydney 2012 clade in the RdRp tree in Figure S1 is shown. While the sequences marked with orange and blue rectangles are monophyletic in the VP1 tree, these sequences cluster apart in the RdRp tree and were therefore acquired in separate recombination events. The RdRp marked with a blue rectangle clusters within the GII.Pe-Sydney 2012 clade, strongly suggesting that this RdRp was acquired from a virus with a Sydney 2012 VP1, indicating cocirculation of Sydney 2012 viruses with the GII.P4 New Orleans 2009-like RdRp and Sydney 2012 viruses with the GII.Pe RdRp. Posterior supports are shown at key nodes

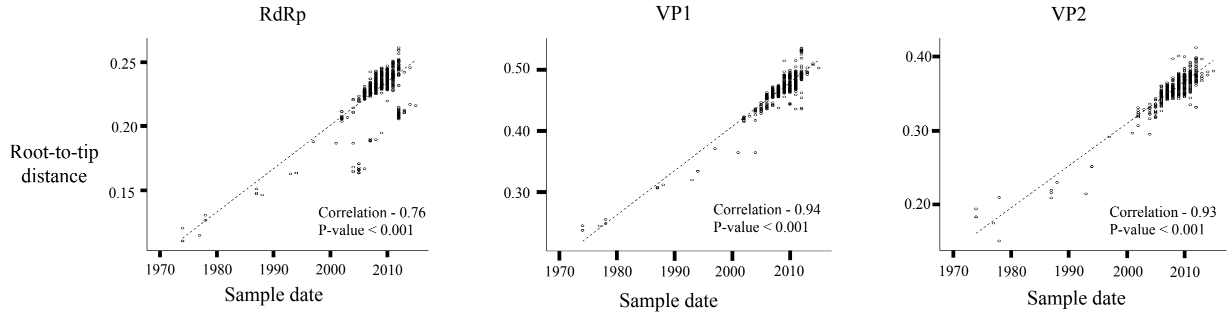


**Fig. S7.**
Temporal evolutionary signal within the RdRp, VP1 and VP2. We reconstructed a nucleotide maximum likelihood tree for each genomic region. Plotted here is the correlation between root-to-tip distance and collection date. The R2 correlation is shown, statistical significance of this correlation was calculated using non-parametric bootstrapping where the collection dates were randomly resampled and the R2 correlation re-calculated 1000 times.

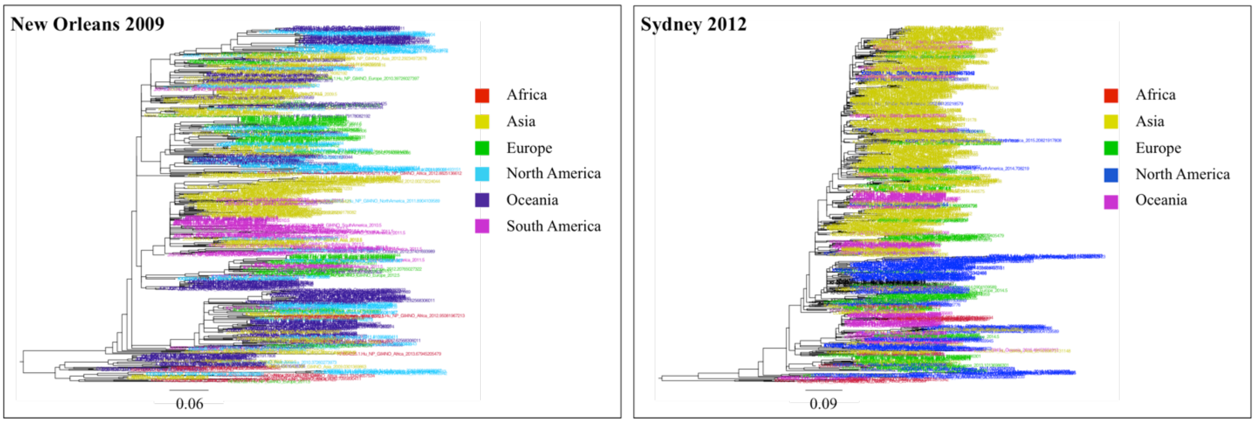


**Fig. S8.**
Interspersion of sequences from each continent within the New Orleans 2009 and Sydney 2012 phylogenetic trees. We reconstructed a nucleotide maximum likelihood tree for the complete New Orleans 2009 and Sydney 2012 VP1 datasets. Each tip label is colored by the continent on which the sequence was collected. The interspersion of sequences from each continent throughout the tree led us to down-sample the sequences from over-represented continents. The scale bar shows the expected number of nucleotide substitutions per site.

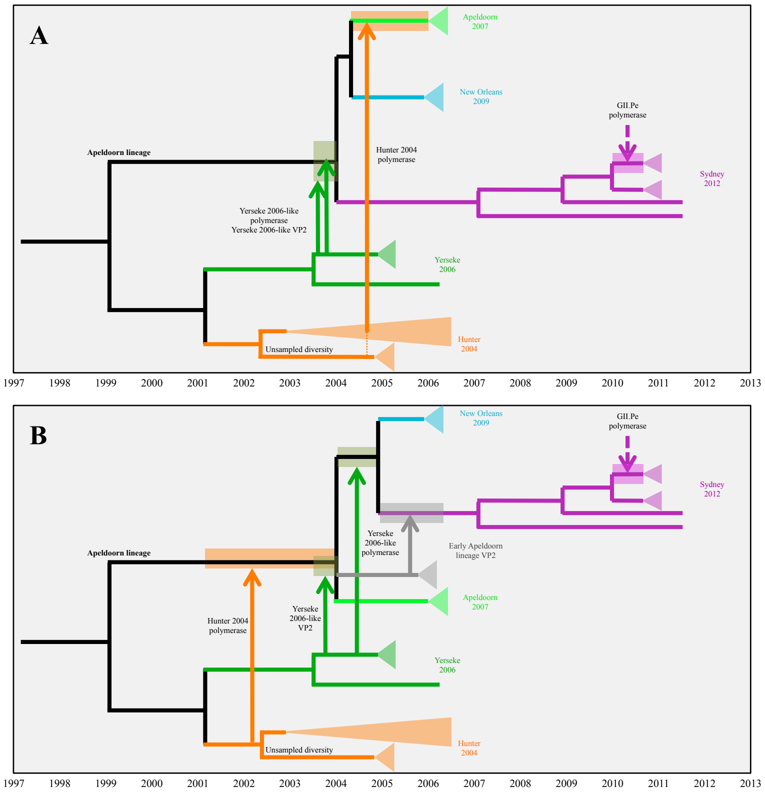


**Fig. S9.**

Plausible scenarios for recombination within the Apeldoorn lineage. Shown are two hypothetical scenarios for the acquisition of RdRp and VP2 genomic regions by the Apeldoorn lineage VP1 (consisting of Apeldoorn 2007, New Orleans 2009 and Sydney 2012) that are consistent with the tree topologies and divergence dates in Figures 1 and S1. The tree backbones are from the VP1 tree in Figure 1 in which the relationship between Apeldoorn 2007, New Orleans 2009 and Sydney 2012 is uncertain. Therefore each scenario depicts a hypothetical relationship between these variants that fits the branching patterns and inferred ancestor dates within the RdRp, VP1 and VP2 phylogenetic trees. Arrows represent a recombination event in which a RdRp or VP2 genomic region is obtained, with the arrow pointing from donor variant to recipient variant. Triangles at the end of branches represent the remaining lineages within the variant. (A) This scenario involves the acquisition of a Yerseke 2006-like VP2 prior to the divergence of the variants within the lineage. The relationship between Apeldoorn 2007, New Orleans 2009 and Sydney 2012 is the same as the well-supported relationship within the VP2 tree, with Sydney 2012 diverging first. The Yerseke 2006 RdRp is acquired prior to divergence of variants within the Apeldoorn lineage and Apeldoorn 2007 acquires a Hunter 2004-like RdRp after its divergence from New Orleans 2009. (B) This scenario involves acquisition of the Yerseke 2006 VP2 and a Hunter 2004-like RdRp prior to divergence of the variants within the Apeldoorn lineage. Here, Apeldoorn 2007 branches first and the Yerseke 2006-like RdRp is acquired leading to the common ancestor of New Orleans 2009 and Sydney 2012. Sydney 2012 acquires its VP2 region from an early Apeldoorn lineage virus that has persisted, explaining the divergence of Sydney 2012 in the VP2 tree. In both scenarios, Sydney 2012 acquires the GII.Pe RdRp after divergence from Apeldoorn 2007 and New Orleans 2009. Scenario B requires one more recombination event than scenario A and requires the persistence of an unsampled early Apeldoorn lineage.

| Genome region | Log10 Bayes factor rejecting strict clock | Evolutionary rate, x$10^{-3}$ substitutions/site/year (95% HPD) | Root date (95% HPD) | GII.P4 RdRp ancestor date (95% HPD) |
|---|---|---|---|---|
| RdRp | 54.89 | 6.67 (5.84, 7.53) | 1941 (1922, 1958) | 1979 (1975-1982) |
| VP1 | 122.97 | 6.83 (6.03, 7.68) | 1943 (1919, 1965) | N/A |
| VP2 | 44.23 | 6.11 (5.35, 6.93) | 1949 (1934, 1963) | N/A |

**Table S1.**
Summary of Bayesian MCMC results. Mean values and 95% HPD intervals were calculated by combining the complete posterior distribution for the parameter of interest from each of the subsampled datasets. The Log10 Bayes factor is the support for rejecting the strict clock model (single substitution rate across the tree) in favor of the relaxed lognormal clock model (each branch on the phylogenetic tree can have a different substitution rate). This Bayes factor therefore suggests variation in the substitution rate along different branches within each dataset.

| GII.4 Variant | US95/96 | Farmington Hills 2002 | Asia 2003 | Hunter 2004 | Yerseke 2006 | Den Haag 2006 | Osaka 2007 | Apeldoorn 2007 | New Orleans 2009 | Sydney 2012 |
|---|---|---|---|---|---|---|---|---|---|---|
| **US95/96** | | 1992 (1989, 1994) | 1992 (1989, 1994) | 1992 (1989, 1994) | 1992 (1989, 1994) | 1992 (1989, 1994) | 1992 (1989, 1994) | 1992 (1989, 1994) | 1992 (1989, 1994) | 1992 (1989, 1994) |
| **Farmington Hills 2002** | | | 1997 (1995, 1999) | 1996 (1994, 1998) | 1996 (1994, 1998) | 1997 (1994, 1999) | 1994 (1991, 1996) | 1996 (1994, 1998) | 1996 (1994, 1998) | 1996 (1994, 1998) |
| Asia 2003 | | | | 1996 (1994, 1998) | 1996 (1994, 1998) | 1996 (1994, 1998) | 1994 (1991, 1996) | 1996 (1994, 1998) | 1996 (1994, 1998) | 1996 (1994, 1998) |
| **Hunter 2004** | | | | | 2001 (1999, 2002) | 1997 (1995, 1999) | 1994 (1991, 1996) | 1999 (1996, 2001) | 1999 (1996, 2001) | 1999 (1996, 2001) |
| Yerseke 2006 | | | | | | 1997 (1995, 1999) | 1994 (1991, 1996) | 1999 (1996, 2001) | 1999 (1996, 2001) | 1999 (1996, 2001) |
| **Den Haag 2006** | | | | | | | 1994 (1991, 1996) | 1997 (1995, 2000) | 1997 (1995, 2000) | 1997 (1995, 2000) |
| Osaka 2007 | | | | | | | | 1994 (1991, 1996) | 1994 (1991, 1996) | 1994 (1991, 1996) |
| Apeldoorn 2007 | | | | | | | | | 2004 (2002, 2005) | 2003 (2002, 2005) |
| **New Orleans 2009** | | | | | | | | | | 2004 (2002, 2005) |

**Table S2.**
Summary of variant divergence times. We calculated the date at which each pair of GII.4 variants diverged in each tree in the VP1 posterior distribution and calculated the mean and 95% HPD of this distribution. Each variant diverges from all of the other sampled variants years prior to pandemic or epidemic emergence, indicating variants evolved independently for years prior to emergence.

| Accession number | Variant | VP1 region present | Country of collection | Age of patient | Reported collection date | Estimated collected date (95% HPD) |
|---|---|---|---|---|---|---|
| EU078408.1 | Farmington Hills 2002 | Complete | USA | NA | 2001 | 2000 (1998-2003) |
| GU937448.1 | Farmington Hills 2002 | P domain | USA | NA | 2000 | 2000 (1998-2003) |
| EU916961.1 | Hunter 2004 | Complete | Tunisia | <12 years | 11-January-2003 | 2004 (2003-2006) |
| EU916960.1 | Hunter 2004 | Complete | Tunisia | <12 years | 11-September-2003 | 2003 (2002-2005) |
| EU916959.1 | Hunter 2004 | Complete | Tunisia | <12 years | 05-April-2003 | 2004 (2003-2006) |
| EU916957.1 | Hunter 2004 | Complete | Tunisia | <12 years | 17-October-2003 | 2004 (2003-2006) |
| EU876890.1 | Hunter 2004 | Complete | France | NA | 2002 | 2004 (2002-2006) |
| GQ845367.2 | New Orleans 2009 | Complete | Australia | NA | November-2008 | 2008 (2006-2010) |
| GQ845345.2 | New Orleans 2009 | Complete | Australia | NA | August-2008 | 2008 (2006-2009) |
| KR131773.1 | New Orleans 2009 | Complete | India | ≤5 years | 2008 | 2009 (2008-2010) |
| KJ735099.1 | Sydney 2012 | Missing first 25 AA | Morocco | NA | 14-September-2011 | 2011 (2009-2012) |
| KF509947.2 | Sydney 2012 | Complete | Canada | NA | September-2011 | 2010 (2007-2012) |
| AB972499.1 | Sydney 2012 | Complete | Japan | NA | 2011 | 2012 (2011-2013) |
| LC005724.1 | Sydney 2012 | Complete | Japan | Nursing home outbreak[a] | December-2011 | 2012 (2010-2014) |
| LC005723.1 | Sydney 2012 | Complete | Japan | Restaurant outbreak[b] | December-2011 | 2011 (2010-2013) |
| LC005722.1 | Sydney 2012 | Complete | Japan | Primary school outbreak[c] | December-2011 | 2011 (2010-2013) |
| LC005721.1 | Sydney 2012 | Complete | Japan | Nursery outbreak[c] | December-2011 | 2011 (2010-2012) |
| LC005720.1 | Sydney 2012 | Complete | Japan | Nursery outbreak[c] | November-2011 | 2012 (2010-2015) |
| KR904236.1 | Sydney 2012 | Complete | South Africa | ≤5 years | 19-December-2011 | 2009 (2006-2012) |
| KR904215.1 | Sydney 2012 | Complete | South Africa | ≤5 years | 06-July-2010 | 2009 (2006-2011) |
| KR904214.1 | Sydney 2012 | Complete | South Africa | ≤5 years | 19-May-2010 | 2009 (2006-2012) |
| KC962462.3 | Sydney 2012 | Complete | South Africa | ≤5 years | November-2011 | 2010 (2007-2013) |
| KF870711.1 | Sydney 2012 | AA 299-end | Spain | NA | February-2011 | 2010 (2008-2015) |
| KF668567.1 | Sydney 2012 | Complete | Italy | Child | 01-November-2011 | 2011 (2010-2013) |
| KF060124.1 | Sydney 2012 | Complete | New Zealand | NA | September-2010 | 2009 (2006-2012) |
| KF060123.1 | Sydney 2012 | Complete | New Zealand | NA | September-2010 | 2009 (2006-2011) |
| KF060122.1 | Sydney 2012 | Complete | New Zealand | NA | September-2010 | 2009 (2006-2012) |
| KX354057.1 | Sydney 2012 | Complete | USA | NA | 26-October-2011 | 2012 (2010-2015) |
| FJ411171.1 | Osaka 2007 | Complete | USA | NA | 2005 | 2003 (1999-2007) |
| EU876884.1 | Osaka 2007 | Complete | Egypt | Between 1 month and 18 years | November-2006 | 2004 (2000-2008) |
| EU876882.1 | Osaka 2007 | Complete | Egypt | Between 1 month and 18 years | September-2006 | 2004 (2000-2008) |

**Table S3.**

Summary of pre-pandemic and pre-epidemic GII.4 sequences. Summary of the 31 sequences with a reported collection date earlier than the year of pandemic/epidemic emergence where the reported collection date was supported by tip dating (see methods). These sequences were taken to be true pre-pandemic/pre-epidemic sequences. While the estimated collection date is here reported to the year, we compared the reported and estimated collection dates to the most precise date possible. Therefore where the reported collection date was given to the nearest day or month, the 95% HPD of the estimated collection date overlaps with that day or month. The proportion of the complete VP1 sequence present in each putative pre-pandemic/pre-epidemic sequence is shown, AA – amino acid.

| Accession number | Variant | VP1 region present | Country of collection | Reported collection date | Estimated collection date (95% HPD) |
|---|---|---|---|---|---|
| KU312299.1 | Farmington Hills 2002 | P2 domain | UK | 1994 | 2002 (2001-2004) |
| EU916958.1 | Hunter 2004 | Missing first 3 AA | Tunisia | 02-January-2003 | 2004 (2003-2005) |
| KU182476.1 | Hunter 2004 | Complete | Tunisia | 2002 | 2004 (2003-2006) |
| KU182477.1 | Hunter 2004 | Complete | Tunisia | 2002 | 2004 (2003-2005) |
| KU182478.1 | Hunter 2004 | Complete | Tunisia | 2002 | 2004 (2003-2006) |
| KU182479.1 | Hunter 2004 | Complete | Tunisia | 2002 | 2004 (2003-2006) |
| KU182480.1 | Hunter 2004 | Complete | Tunisia | 2002 | 2004 (2003-2006) |
| KU182481.1 | Hunter 2004 | Complete | Tunisia | 2002 | 2004 (2003-2006) |
| KU182482.1 | Hunter 2004 | Complete | Tunisia | 2002 | 2004 (2003-2006) |
| KU182483.1 | Hunter 2004 | Complete | Tunisia | 2002 | 2004 (2003-2006) |
| KU312301.1 | Hunter 2004 | P2 domain | UK | 1995 | 2004 (2002-2007) |
| KU312302.1 | Hunter 2004 | P2 domain | UK | 1995 | 2004 (2002-2006) |
| KU312303.1 | Hunter 2004 | P2 domain | UK | 1995 | 2004 (2002-2007) |
| AB972505.1 | Sydney 2012 | Complete | Japan | 2011 | 2013 (2012-2014) |
| AB972504.1 | Sydney 2012 | Complete | Japan | 2011 | 2013 (2012-2014) |
| AB972503.1 | Sydney 2012 | Complete | Japan | 2011 | 2013 (2012-2015) |
| AB972502.1 | Sydney 2012 | Complete | Japan | 2011 | 2013 (2012-2014) |
| KU312300.1 | Sydney 2012 | P2 domain | UK | 1994 | 2013 (2011-2015) |
| FJ411172.1 | Osaka 2007 | Complete | USA | 2006 | 2001 (1997-2004) |

**Table S4.**

Summary of the 19 sequences with a reported collection date earlier than the year of pandemic/epidemic emergence where the reported collection date was not supported by tip dating (see methods). These sequences were not taken to be true pre-pandemic/pre-epidemic sequences. Possible reasons for this discrepancy include mis-reporting of the collection date, sample contamination, sample mis-labelling and inaccurate estimation of the collection date. The 95% HPD of the estimated collection date of sequence EU916958.1 does not overlap with the reported collection date to the level of the day. AA – amino acid.

| GII.4 Variant | Common ancestor date | | |
|---|---|---|---|
| | **RdRp (95% HPD)** | **VP1 (95% HPD)** | **VP2 (95% HPD)** |
| **US95/96** | December 1993 (November 1991-January 1996) | December 1993 (June 1991-April 1996) | April 1995 (September 1993-September 1996) |
| **Farmington Hills 2002** | July 2000 (August 1999-April 2001) | December 2000 (February 2000-September 2001) | September 2000 (October 1999-July 2001) |
| Asia 2003 | August 2002 (June 2001-August 2003) | February 2001 (May 1999-November 2002) | March 2002 (November 2000-April 2003) |
| **Hunter 2004** | September 2002 (August 2001-August 2003) | December 2002 (March 2002-September 2003) | April 2002 (September 2000-August 2003) |
| Yerseke 2006 | June 2002 (January 2001-October 2003) | July 2003 (April 2002-October 2004) | April 2003 (April 2002-April 2004) |
| **Den Haag 2006** | March 2004 (February 2003-April 2005) | February 2004 (October 2002-April 2005) | January 2004 (October 2002-April 2005) |
| Osaka 2007 | February 2006 (January 2005-January 2007) | September 2005 (June 2004-November 2006) | December 2005 (November 2004-November 2006) |
| Apeldoorn 2007 | April 2005 (April 2004-March 2006) | January 2006 (December 2004-December 2006) | April 2006 (June 2005-January 2007) |
| **New Orleans 2009** | April 2005 (December 2003-July 2006) | December 2005 (October 2004-February 2007) | October 2006 (November 2005-August 2007) |
| **Sydney 2012** | October 2009 (August 2008-September 2010) | February 2007 (April 2005-October 2008) | May 2006 (August 2004-April 2008) |

**Table S5.**
The common ancestor date of each GII.4 variant is shown for the RdRp, VP1 and VP2. This was estimated by combining the posterior distribution of the common ancestor date for each subsampled dataset. The common ancestor of the Sydney 2012 RdRp is the common ancestor of the GII.Pe RdRps found with the Sydney 2012 VP1.

| Variant | Nonstructural polyprotein | VP1 | VP2 |
|---|---|---|---|
| Farmington Hills 2002 | P44S, I52T, M246V, R278K, V525I, T650A, K729R, A782T, N787S, K807R, A850V, S1270N, K1552R | *S9N*, D298N, K329R, S355D, V365I, T368N, S394G, N407S, A534T | N23S, K80E, Q83R, A97S, S155F, T158V |
| Hunter 2004 | Y759H, S760N, *K807R* | Node 1 - H297Q, N372S, K382R, N412D, T425S<br>Node 2 - N9T, N407D, G413S, S425T, A465S | I101V, L140S, S162A |
| Den Haag 2006 | G27K, V28M, L29F, I79V, V85A, A104T, A261T, L283I, Y327F, A336V, V525I, T650A, R738K, A782T, T791A, V830I, M833V, D983E, S1182G, R1420K, N1575D, T1618S, A1642T | T15A, P174S, H297R, Q306L, R339K, S352Y, V356A, H357P, N372E, G378H, N407S, G413V | V33I, E34D, T144I, A148T, T149P, V150T, P159S, V168I, S187N, K262R |
| New Orleans 2009 | *V525I*, I750V, *V779I*, K1210T, *S1610P*, E1613G | T294A, A340T, A359S | *H130R*, T164I |
| Sydney 2012 | Circulated with multiple nonstructural polyprotein genotypes | $\text{Sydney}^{\text{Anc}}_{\text{All}}$ – T15A<br>$\text{Sydney}^{\text{Anc}}_{\text{Pand}}$- *H297R*, S310N, A368E, N373H, *N393S*, *A395T*, *A539V*, *L540V* | $\text{Sydney}^{\text{Anc}}_{\text{All}}$- A148D<br>$\text{Sydney}^{\text{Anc}}_{\text{Pand}}$- T158K, N205S |

**Table S6.**
Summary of nonsynonymous substitutions leading to the five most recent pandemic GII.4 variants. We used ancestral reconstruction to identify the nonsynonymous substitutions that occurred along the phylogenetic branch leading to the common ancestor of each of the five most recent pandemic GII.4 variants within each genomic region. These substitutions delimit that variant from the other, typically unsampled, lineages present at the time of pandemic emergence and so are likely to encode the characteristic(s) that enabled pandemic emergence. Substitutions in italics are unlikely to have been important for pandemic emergence, as the residue(s) at these sites was the same within the preceding pandemic variant. Red substitutions occurred at VP1 sites within known blockade epitopes (Lindesmith et al. 2012). There are two potential common ancestor nodes for Hunter 2004; the substitutions leading to each of these nodes are shown here.

| VP1 variant | RdRp or VP2 acquired | Inferred date of recombination event (95% HPD) |
|---|---|---|
| Asia 2003 | GII.P12 RdRp | March 1999 (March 1996-October 2001) |
| Osaka 2007 | GII.Pe RdRp | June 2000 (October 1994-January 2006) |
| Osaka 2007 | Den Haag 2006-like VP2 | March 2005 (December 2003-August 2006) |
| Apeldoorn lineage | Yerseke 2006-like VP2 | March 2004 (February 2003-April 2005) |
| Apeldoorn lineage | Yerseke 2006-like RdRp | July 2004 (January 2003-November 2005) |
| Sydney 2012 | GII.Pe RdRp | May 2010 (July 2009-February 2011)<br>January 2012 (July 2011-May 2012)<br>February 2012 (August 2011-July 2012) |

**Table S7.**
Summary of recombination events and dates in the GII.4 lineage. Shown are the dates at which the GII.4 VP1 variants acquired a new RdRp or VP2. Recombination events were initially identified on the basis of well-supported topological differences between the VP1 tree and the RdRp or VP2 tree. The date of the recombination event was calculated from the posterior distribution of trees from the corresponding dataset by identifying the branch along which the recombination event occurred in each tree. Sydney 2012 acquired the GII.Pe RdRp in at least three independent recombination events. Additional recombination events likely occurred in the Apeldoorn lineage (which consists of the Apeldoorn 2007, New Orleans 2009 and Sydney 2012 variants). Further information on recombination within the Apeldoorn lineage is shown in Figures S6 and S9.

| Most likely recombination breakpoint | GII.4 variant 5′ to breakpoint | GII.4 variant 3′ to breakpoint | Number of sequences | Accession numbers |
|---|---|---|---|---|
| 537 | Den Haag 2006 | New Orleans 2009 | 12 | KF712501.1, KF429790.1, KF712491.1, KF429762.1, KF712498.1, KF712505.1, KF712495.1, KF429785.1, KF429776.1, JX459900.1, KF712502.1, AB933738.1 |
| 537 | Den Haag 2006 | Apeldoorn 2007 | 1 | AB541362.1 |
| 537 | New Orleans 2009 | Apeldoorn 2007 | 1 | JX448566.1 |
| 314 | Den Haag 2006 | Apeldoorn 2007 | 1 | KF712492.1 |
| 314 | New Orleans 2009 | Den Haag 2006 | 3 | KF196287.1, AB933682.1, AB447434.1 |
| 314 | Unclear | Osaka 2007 | 1 | GQ845368.2 |

**Table S8.**
Summary of putative recombination events within the GII.4 VP1. We identified putative recombination events using SBP. The most likely recombination breakpoint was defined as the position with the greatest gain in Akaike information criteria (AIC) when splitting the alignment on that alignment site compared with the null model. This breakpoint is therefore approximate. The number of sequences that cluster differently on either side of the putative breakpoint but cluster together on both sides of the putative breakpoint is shown with the variant within which the sequences cluster. Removal of the sequences in this table resulted in loss of the recombination signal.

| Variant name | Number of RdRp sequences | Number of VP1 and VP2 sequences | Subsampled |
|---|---|---|---|
| GII.4 1970s (GII.P1 RdRp) | 6 | 6 | No |
| Bristol 1993 | 2 | 2 | No |
| Camberwell 1994 | 5 | 5 | No |
| US95/96 | 3 | 3 | No |
| Lanzhou 2001 | 1 | 1 | No |
| Farmington Hills 2002 | 18 | 18 | No |
| Asia 2003 (GII.P12 RdRp) | 15 | 15 | No |
| Hunter 2004 | 17 | 17 | No |
| Yerseke 2006 | 12 | 12 | No |
| Den Haag 2006 | 551 | 559 | Yes, 41 sequences |
| Osaka 2007 (GII.Pe RdRp) | 5 | 5 | No |
| Apeldoorn 2007 | 37 | 31 | No |
| New Orleans 2009 | 141 | 134 | Yes, 41 sequences |
| Sydney 2012 (GII.Pe RdRp) | 36 | 41 | No |
| GII.4 could not assign strain | 3 | 3 | No |

**Table S9.**
Summary of the GII.4 sequences included in the dataset used to reconstruct the temporal history of the complete GII.4 genotype. The number of sequences from each GII.4 variant is shown following the removal of potentially recombinant sequences. The RdRp genotype is shown in parentheses for each GII.4 variant that is not found with the GII.P4 RdRp. Den Haag 2006 and New Orleans 2009 were randomly subsampled to the number of sequences present in the third most prevalent variant. Sydney 2012 and Osaka 2007 are found with the GII.Pe RdRp, therefore there are 41 GII.Pe sequences in the RdRp dataset.

| GII.4 Variant | Polymerase ancestor date (95% HPD) | | | VP1 ancestor date (95% HPD) | | | VP2 ancestor date (95% HPD) | | |
|---|---|---|---|---|---|---|---|---|---|
| | Sample 1 | Sample 2 | Sample 3 | Sample 1 | Sample 2 | Sample 3 | Sample 1 | Sample 2 | Sample 3 |
| **US95/96** | 1993 (1991,1995) | 1993 (1991,1996) | 1994 (1992,1996) | 1994 (1991,1996) | 1993 (1991,1996) | 1993 (1991,1996) | 1994 (1993,1996) | 1994 (1993,1996) | 1994 (1993,1996) |
| **Farmington Hills 2002** | 2000 (1999,2001) | 2000 (1999,2001) | 2000 (1999,2001) | 2001 (2000,2001) | 2000 (1999,2001) | 2001 (2000,2001) | 1999 (1998,2000) | 1999 (1998,2000) | 2000 (1998,2000) |
| Asia 2003 | 2002 (2001,2003) | 2002 (2001,2003) | 2002 (2001,2003) | 2001 (1999,2002) | 2001 (1999,2002) | 2001 (1999,2002) | 2001 (2000,2002) | 2001 (1999,2002) | 2001 (2000,2002) |
| **Hunter 2004** | 2002 (2001,2003) | 2002 (2001,2003) | 2002 (2001,2003) | 2003 (2002,2003) | 2002 (2002,2003) | 2002 (2002,2003) | 2002 (2001,2003) | 2002 (2001,2003) | 2002 (2001,2003) |
| Yerseke 2006 | 2002 (2001,2003) | 2002 (2001,2003) | 2002 (2001,2003) | 2003 (2002,2004) | 2003 (2002,2004) | 2003 (2002,2004) | 2002 (2001,2003) | 2002 (2001,2003) | 2002 (2002,2004) |
| **Den Haag 2006** | 2004 (2003,2005) | 2004 (2003,2005) | 2003 (2002,2004) | 2004 (2002,2005) | 2004 (2002,2005) | 2004 (2003,2005) | 2002 (2001,2003) | 2002 (2001,2004) | 2003 (2002,2004) |
| Osaka 2007 | 2006 (2005,2007) | 2006 (2005,2007) | 2006 (2005,2006) | 2005 (2004,2006) | 2005 (2004,2006) | 2005 (2004,2006) | 2005 (2004,2006) | 2005 (2003,2006) | 2005 (2004,2006) |
| Apeldoorn 2007 | 2005 (2004,2006) | 2005 (2004,2006) | 2005 (2004,2006) | 2006 (2005,2007) | 2005 (2004,2006) | 2005 (2004,2006) | 2005 (2004,2006) | 2005 (2004,2006) | 2005 (2004,2006) |
| **New Orleans 2009** | 2005 (2003,2006) | 2005 (2004,2006) | 2005 (2003,2006) | 2006 (2004,2007) | 2005 (2004,2007) | 2005 (2004,2006) | 2006 (2005,2006) | 2005 (2004,2006) | 2005 (2005,2006) |
| **Sydney 2012** | 2009 (2008,2010) | 2009 (2008,2010) | 2009 (2008,2010) | 2007 (2005,2008) | 2007 (2005,2008) | 2006 (2005,2008) | 2005 (2003,2007) | 2005 (2003,2007) | 2005 (2003,2007) |

**Table S10.**
Comparison of variant ancestor dates in each subsampled dataset. Samples 1, 2 and 3 contain a different random subsample of 41 sequences from Den Haag 2006 and 41 sequences from New Orleans 2009. The Sydney 2012 RdRp ancestor date is the common ancestor of the GII.Pe RdRps found with the GII.4 Sydney 2012 VP1.

| Country | Number of New Orleans 2009 sequences | Number of Sydney 2012 sequences | Continent |
|---|---|---|---|
| Albania | 1 | 0 | Europe |
| Australia | 144 | 56 | Oceania |
| Bangladesh | 8 | 4 | Asia |
| Brazil | 65 | 0 | South America |
| Canada | 6 | 6 | North America |
| China | 21 | 234 | Asia |
| Denmark | 15 | 23 | Europe |
| France | 1 | 7 | Europe |
| Germany | 0 | 11 | Europe |
| Hungary | 1 | 0 | Europe |
| India | 10 | 3 | Asia |
| Italy | 10 | 4 | Europe |
| Japan | 44 | 87 | Asia |
| Morocco | 0 | 1 | Africa |
| Netherlands | 2 | 2 | Europe |
| New Zealand | 0 | 32 | Oceania |
| Russia | 8 | 2 | Europe |
| Singapore | 16 | 0 | Asia |
| Slovenia | 0 | 56 | Europe |
| South Africa | 18 | 16 | Africa |
| South Korea | 29 | 6 | Asia |
| Spain | 5 | 3 | Europe |
| Sweden | 12 | 0 | Europe |
| Taiwan | 16 | 27 | Asia |
| UK | 19 | 0 | Europe |
| USA | 89 | 109 | North America |
| Vietnam | 26 | 7 | Asia |

| Continent | Number of New Orleans 2009 sequences | Number of Sydney 2012 sequences |
|---|---|---|
| Africa | 18 | 17 |
| Asia | 170 | 368 |
| Europe | 74 | 108 |
| North America | 95 | 115 |
| Oceania | 144 | 88 |
| South America | 65 | 0 |

**Table S11.**

Summary of the collection countries and continents of sequences used in phylogeographic analysis. The number of sequences from each country and from each continent is shown for the New Orleans 2009 and Sydney 2012 VP1 datasets. Sequences from Asia and Oceania were randomly subsampled in the New Orleans 2009 dataset and sequences from Asia were randomly subsampled in the Sydney 2012 dataset.

| GII.4 variant | Number of VP1 sequences | Number of VP2 sequences |
|---|---|---|
| CHDC 1970s | 8 | 8 |
| Tokyo 1980s | 2 | 2 |
| Bristol 1993 | 3 | 3 |
| Camberwell 1994 | 7 | 5 |
| US95/96 | 79 | 5 |
| Farmington Hills 2002 | 92 | 61 |
| Lanzhou 2002 | 11 | 2 |
| Asia 2003 | 42 | 16 |
| Kaiso 2003 | 5 | 0 |
| Hunter 2004 | 59 | 17 |
| Den Haag 2006 | 858 | 629 |
| Yerseke 2006 | 46 | 18 |
| Apeldoorn 2007 | 69 | 36 |
| Osaka 2007 | 30 | 4 |
| New Orleans 2009 | 396 | 158 |
| Sydney 2012 | 480 | 157 |
| GII.4 could not assign strain | 22 | 6 |

**Table S12.**
Summary of the datasets used in reconstruction of the Sydney 2012 ancestral VP1 sequences and to identify the nonsynonymous substitutions that occurred in VP1 and VP2 leading to each pandemic GII.4 variant. Strain typing was carried out using the norovirus genotyping tool. All variant assignments were confirmed by subsequent phylogenetic analyses. Variant names used are those returned by the norovirus genotyping tool for VP1.

| Genotype | Number of samples |
|---|---|
| GII.P1 | 6 |
| GII.P2 | 1 |
| GII.P3 | 5 |
| GII.P4 Bristol 1993 | 2 |
| GII.P4 Camberwell 1994 | 5 |
| GII.P4 US95/96 | 5 |
| GII.P4 Farmington Hills 2002 | 18 |
| GII.P4 Lanzhou 2002 | 1 |
| GII.P4 Hunter 2004 | 16 |
| GII.P4 Den Haag 2006 | 595 |
| GII.P4 Yerseke 2006 | 11 |
| GII.P4 Apeldoorn 2007 | 43 |
| GII.P4 Osaka 2007 | 1 |
| GII.P4 New Orleans 2009 | 150 |
| GII.P4 Could not assign strain | 4 |
| GII.P5 | 1 |
| GII.P6 | 3 |
| GII.P7 | 42 |
| GII.P8 | 2 |
| GII.P11 | 1 |
| GII.P12 | 36 |
| GII.P13 | 1 |
| GII.P15 | 1 |
| GII.P16 | 9 |
| GII.P17 | 56 |
| GII.P18 | 1 |
| GII.P20 | 1 |
| GII.P21 | 30 |
| GII.P22 | 17 |
| GII.Pa | 2 |
| GII.Pc | 3 |
| GII.Pe | 85 |
| GII.Pg | 17 |
| GII.Pj | 1 |
| GII.Pm | 2 |
| GII.Pp | 1 |
| GII could not assign genotype | 4 |
| GIV | 4 |

**Table S13.**
Summary of the dataset used in reconstruction of the nonsynonymous substitutions that occurred leading to each pandemic GII.4 variant within the nonstructural polyprotein. Strain typing was carried out using the norovirus genotyping tool. All variant assignments were confirmed by subsequent phylogenetic analyses. Variant names are those returned by the norovirus genotyping tool.
